# Topological defects in the nematic order of actin fibers as organization centers of *Hydra* morphogenesis

**DOI:** 10.1101/2020.03.02.972539

**Authors:** Yonit Maroudas-Sacks, Liora Garion, Lital Shani-Zerbib, Anton Livshits, Erez Braun, Kinneret Keren

## Abstract

Animal morphogenesis arises from the complex interplay between multiple mechanical and biochemical processes with mutual feedback. Developing an effective, coarse-grained description of morphogenesis is essential for understanding how these processes are coordinated across scales to form robust, functional outcomes. Here we show that the nematic order of the supra-cellular actin fibers in regenerating *Hydra* defines a slowly-varying field, whose dynamics provide an effective description of the morphogenesis process. We show that topological defects in this field, which are long-lived yet display rich dynamics, act as organization centers with morphological features developing at defect sites. These observations suggest that the nematic orientation field can be considered a “mechanical morphogen” whose dynamics, in conjugation with various biochemical and mechanical signaling processes, result in the robust emergence of functional patterns during morphogenesis.

---

Animal morphogenesis involves multiple mechanical and biochemical processes, spanning several orders of magnitude in space and time, from local dynamics at the molecular level to global, organism-scale morphology. How these numerous processes are coordinated and integrated across scales to form robust, functional outcomes remains an outstanding question ^1–4^. Developing an effective, coarse-grained description of morphogenesis can provide essential insights towards addressing this important challenge. Here, we focus on whole-body regeneration in *Hydra*, a small fresh-water predatory animal, and provide an effective description of the morphogenesis process that is based on the dynamic organization of the supra-cellular actin fibers in regenerating tissues ^5, 6^.

*Hydra* is a classic model system for morphogenesis, thanks to its simple body plan and remarkable regeneration capabilities. Historically, research on *Hydra* regeneration inspired the development of many of the fundamental concepts on the biochemical basis of morphogenesis, including the role of an “organizer” ^7^, the idea of pattern formation by reaction-diffusion dynamics of morphogens ^8, 9^, and the concept of positional information ^10^. However, in these studies, as well as in the majority of subsequent works, the role of mechanics in *Hydra* morphogenesis was largely overlooked. Here we revisit this classic model system, and investigate the cytoskeletal dynamics during the regeneration process from a biophysical point of view, revealing a previously unappreciated relation between cytoskeletal organization and the morphogenesis process.

A mature *Hydra* has a simple uniaxial body plan, with a head on one end and a foot on the other. Its tubular body consists of a double layer of epithelial cells, which contain highly organized arrays of parallel supra-cellular actin fibers (Fig. S1). The fibers are globally aligned along the body axis in the outer (ectoderm) layer, and perpendicular to the body axis in the inner (endoderm) layer ^5^. The actin fibers are decorated with myosin motors, forming contractile bundles called myonemes, which are akin to muscular structures in other organisms. The fibers lie along the basal surfaces of each epithelial layer and are connected through cell-cell junctions to form supra-cellular bundles, which exhibit long range directional order over scales comparable to the size of the animal, ranging from ~300 μm in small regenerated *Hydra* to several mm in fully grown *Hydra*. The myonemes are coupled via adhesion complexes to a thin viscoelastic extracellular matrix layer (the mesoglea) sandwiched between the two cell layers (Fig. S1) ^11^.

Large-scale supra-cellular arrays of contractile actin-myosin fibers are a common organizational theme found in many animal tissues ^12–14^. While the mechanism for the development of these parallel fiber arrays is not entirely clear, mechanical feedback has been shown to play a central role in their formation and alignment ^14, 15^. Such parallel alignment is characteristic of systems with nematic order, i.e. systems in which the microscopic constituents tend to locally align parallel to each other exhibiting long-range orientational order ^16^. Nematic systems can be described by a director field which denotes the local orientation, and its spatio-temporal evolution reflects the dynamics of the system. Such systems can be further classified as “active” if their constituents consume energy and generate forces, giving rise to a wealth of interesting dynamic behaviors ^17^. Recently, the framework of active nematics has been found to be informative in understanding various phenomena in a variety of biophysical systems ^17, 18^ including *in vitro* cytoskeletal networks ^19–23^, cell monolayers ^24–26^ and bacterial cultures ^27^. Here we describe the organization of the supra-cellular actin fibers in regenerating *Hydra* as an active nematic system.

*Hydra* contains two distinct perpendicular arrays of nematic supra-cellular actin fibers in the ectoderm and in the endoderm (Fig. S1). Here we primarily consider the thicker and more continuous ectodermal actin fibers, which are the most prominent cytoskeletal feature affecting *Hydra* mechanics ^5, 6^. The director field describing the alignment of the ectodermal fibers defines a slowly-varying field that persists over extended spatial and temporal scales compared to molecular and cellular scales (Fig. S2). In particular, unlike other tissue characteristics (e.g. cell shape anisotropy, local curvature), the fiber orientation is maintained even under extensive tissue deformations. As such, the fiber orientation field provides a valuable coarse-grained description of the system. In that respect, the aligned fibers serve a dual role: an integrated probe reporting on a host of underlying biochemical and mechanical processes, and the major force generator in the system. Importantly, the contractile actin fibers simultaneously contribute and respond to the stresses and deformations in the tissue, consuming energy and maintaining the system far from thermal equilibrium. Such coupling between a nematic field and surface deformations in non-equilibrium systems introduces rich physics, which are only beginning to be explored experimentally ^20, 23^ and theoretically ^28, 29^. Note that unlike many of the recently studied active nematic systems ^17^, here the fibers do not propel themselves, but rather generate active stresses upon contraction.

One of the key features of nematic systems is the possible existence of singularities in the director field (locally disordered regions), which are called topological defects, and are classified by the number of rotations of the nematic director around the singular point (called topological charge; see Methods) ^16^. The total charge of defects in a closed system is constrained by topology ^16^. In particular, for a nematic on the surface of a closed shell, which is the relevant case for *Hydra*, the total charge has to equal +2. Interestingly, topological defects in the nematic order have been shown to induce local changes in the dynamics of cell monolayers *in vitro* ^24, 26^, that were related to mechanical feedback whereby alteration of the local stress field near defects triggered changes in cellular behavior (e.g. cell death and extrusion in epithelial sheets ^24^).

Here we follow the nematic dynamics of the actin fibers during *Hydra* regeneration from excised tissue pieces by live microscopy. The fiber organization in excised tissues, which is inherited from the parent animal, is only partially maintained as the tissues fold into spheroids ^6^. During regeneration, the regions with aligned fibers induce order in neighboring regions, yet nematic topological defects in the actin fiber alignment must appear, once the nematic order extends throughout the system, due to the topological constraint. We show that these topological defects are long-lived, and display rich dynamics, including motility, as well as merging and annihilation of defect pairs. Importantly, we find that the dynamics of defects during regeneration are closely related to the formation of the main functional morphological features, namely the head and the foot, with the fiber alignment ultimately determining the regenerated body axis. The observed relation between defect dynamics and the regeneration process offers a complementary biophysical viewpoint on morphogenesis, with the nematic orientation field serving as a “mechanical morphogen” and sites of defects in this field acting as organization centers of the developing body plan.

## Nematic organization of the supra-cellular actin fibers in mature Hydra

Mature *Hydra* have a parallel array of actin fibers in the ectoderm which is aligned with their body axis and can be described by a nematic director field (Fig. 1A,B). Since the *Hydra* body, where the fibers reside, forms a closed 2D hollow shell, the topological constraint on the net charge implies that topological defects are unavoidable. Importantly, the topological defects in mature *Hydra* coincide with the morphological features of the animal. Specifically, two +1 point defects are found at the apex of the head and foot regions; a +1 defect at the tip of the mouth (hyptosome; Fig. 1C), and a +1 defect at the base of the foot, surrounding the basal disc (Fig. 1D). The body axis of the animal is aligned between these two defects, which have a total charge of +2, equal to the topologically required charge. Additionally, mature *Hydra* have a variable number of tentacles that contain further topological defects; each tentacle has a +1 defect at its tip (Fig. 1E) that is accompanied by two −1/2 defects at opposite sides of its base (Fig. 1F). The total defect charge of a tentacle is thus equal to 0, and hence does not contribute to the net defect charge in the animal. Similarly, multi-axis *Hydra* (Fig. S3) or *Hydra* with an emerging bud ^5^, contain an additional +1 defect at the extra head/foot, which is balanced by two −1/2 defects on either side of the junction between the two body axes (Fig. S3C). Thus, while the constraint on the topological charge necessitates the presence of defects and restricts the possible defect configurations to have a net charge equal to +2, it does not prescribe the body plan of the regenerated animal since the presence of additional structures with opposite-charged defects that cancel each other is not precluded. In general, we find that +1 defects are localized in regions of high positive surface curvature, whereas −1/2 defects reside in saddle regions with negative curvature. Interestingly, mature *Hydra* contain only +1 defects and −1/2 defects, whereas regenerating tissue spheroids exhibit +1/2 defects, in addition to +1 and −1/2 defects (Fig. 1G-I). The same pattern and type of defects are observed in the inner endoderm layer (Fig. S4). Note that since the orientation of the supra-cellular actin fibers in the endoderm is perpendicular to the ectodermal fibers, the +1 defects in the endoderm are “vortex”-shaped ^30^, rather than the “aster”-shaped defects observed in the ectoderm (Fig. 1C,G).

**Fig. 1.**
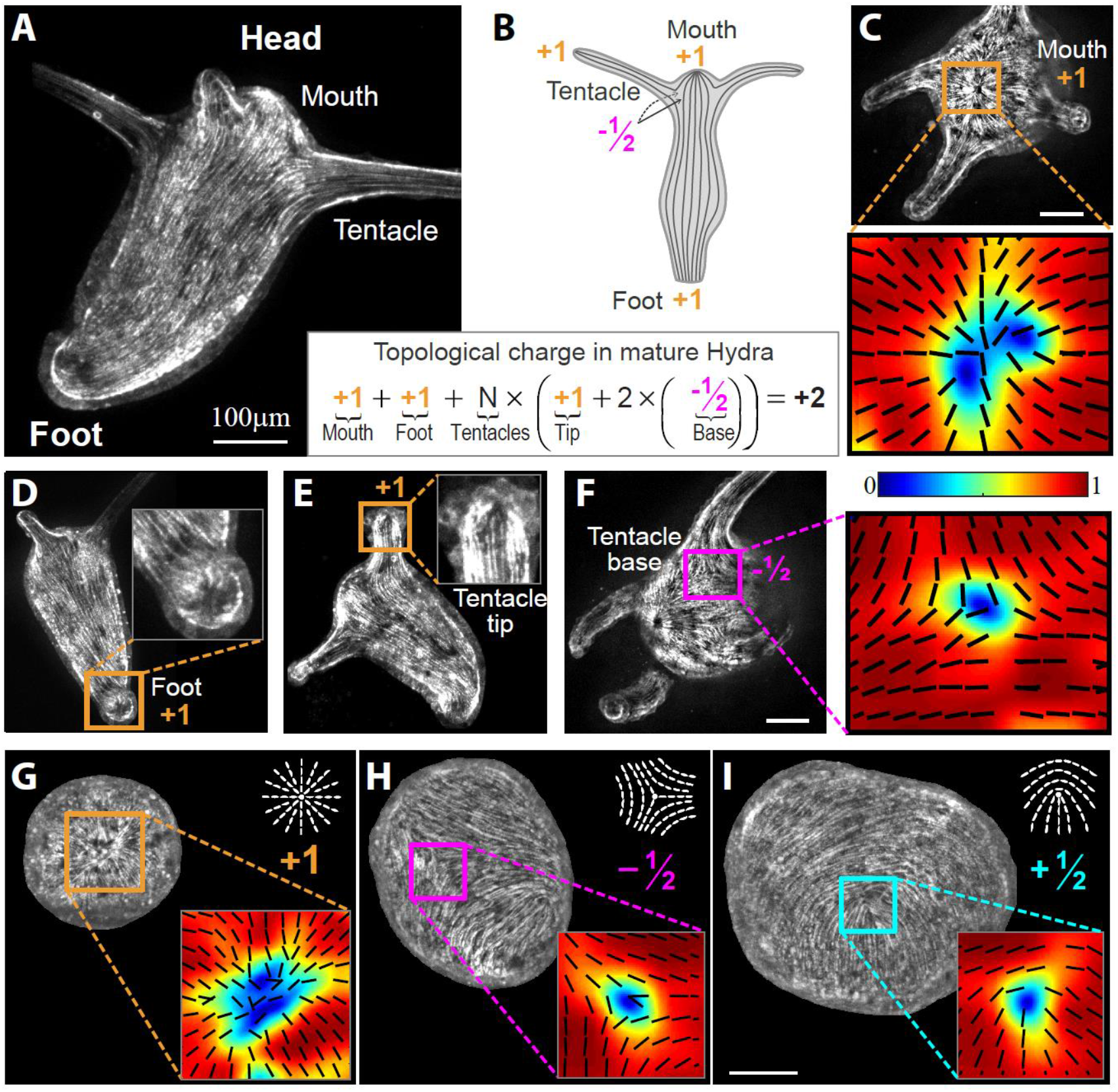
Topological defects in *Hydra*. (A) Image showing the nematic actin fiber organization in the ectoderm of a small mature *Hydra*. (B) Schematic illustration of the nematic actin fiber organization in a mature *Hydra*. The topological defects in the nematic organization coincide with the morphological features of the animal, with defects localized at the mouth (+1), foot (+1) and tentacles (+1 at the tip, and two −1/2 at the base). The sum of defect charge is constrained by topology to be equal to +2. (C-F) Images of the actin fiber organization containing topological defects localized at the functional morphological features of a mature *Hydra*: the tip of the head (C), the apex of the foot (D), and the tip (E) and base (F), respectively, of a tentacle. The zoomed maps in (C, F) depict the actin fiber orientation extracted from the image intensity gradients ^36^ (black lines) and the local order parameter (color; see Methods). (G-I) Examples of the types of topological defects found in regenerating tissue spheroids imaged in different samples: +1 defect (G), −1/2 defect (H) and +1/2 defect (I). Inset: zoomed maps of the corresponding actin fiber orientation and the local order parameter. All images shown are 2D projections extracted from 3D spinning-disk confocal z-stacks of transgenic *Hydra* expressing lifeact-GFP in the ectoderm.

## *Nematic organization of the supra-cellular actin fibers in regenerating Hydra* tissues

How does the nematic configuration found in mature *Hydra* arise during regeneration from a tissue segment? To address this question, we followed actin dynamics in regenerating tissue fragments excised from transgenic *Hydra* expressing lifeact-GFP in the ectoderm (Fig. 2). Initially, an excised tissue fragment is characterized by a perfect nematic array of actin fibers, inherited from the parent animal, with no defects (Fig. S5A) ^6^. The excised tissue folds within a couple of hours, in an active actomyosin-dependent process, to form a closed spheroid ^6^. The change in surface topology, from a tissue fragment with open boundaries to a closed, hollow spheroid, necessarily implies that the nematic order cannot be maintained throughout the system. Indeed, the folding process is accompanied by a dramatic reorganization during which actin fibers near the center of the fragment are stably maintained, whereas fibers near the closure region of the spheroid undergo extensive remodeling and the nematic organization is essentially lost there ^6^. We find that the ordered nematic fiber array occupies roughly half of the spheroid’s surface area, while the remaining surface is essentially devoid of aligned fibers (Fig. 2B, left).

**Fig. 2.**
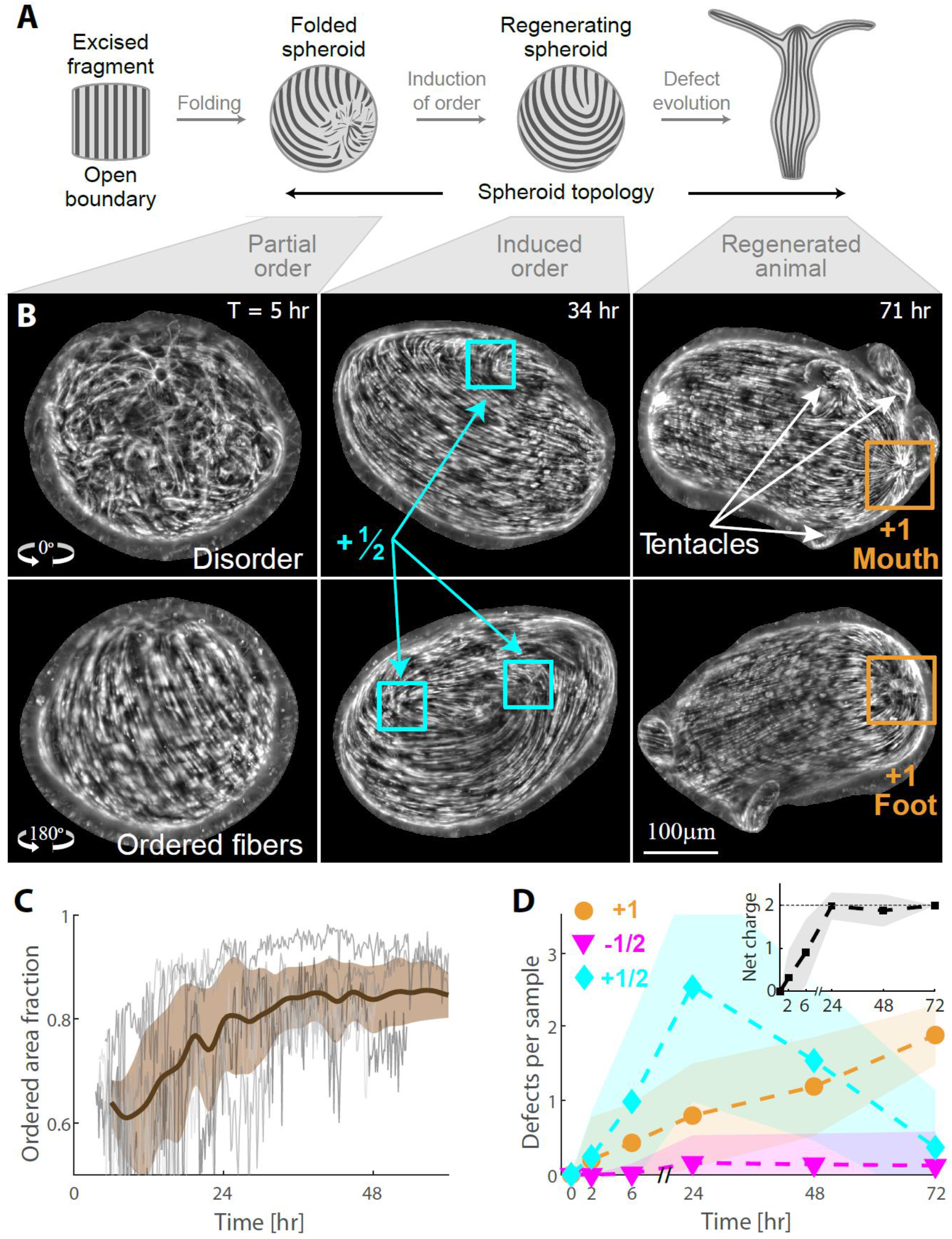
Nematic actin organization in regenerating tissue fragments. (A) Schematic illustration of the nematic organization of the actin fibers at different stages during the regeneration of an excised tissue fragment. Immediately after excision (left), the tissue fragment contains an ordered array of fibers. After ~2 hours (middle-left), the tissue is folded into a closed spheroid with partial nematic order and an extended disordered region. Induction of order over time, leads to the formation of localized topological defects (middle-right), which evolve over the course of a couple of days as the tissue regenerates into a mature *Hydra* (right). (B) Images from a time-lapse light-sheet movie of a regenerating tissue fragment (Movie 1). At each time-point, projections from two opposite sides depict the actin fiber organization in the regenerating tissue. At *T*=5 hr, half of the spheroid is disordered (top), whereas a nematic array of fibers is observed on the other half (bottom). At *T*=34 hr, ordered fibers are apparent everywhere except for localized point defects. Three +1/2 defects are visible in the images. At *T*=71 hr, the defect configuration of a mature *Hydra* is apparent with two +1 defects at opposite sides of the tissue marking the tips of the head and the foot, respectively, and additional defects in the emerging tentacles. (C) The fraction of area containing ordered fibers in regenerating fragments as a function of time (see Methods). The graph depicts the ordered area fraction for 5 different regenerating fragments that were imaged by light-sheet microscopy from 4 directions (thin lines), together with the average trend (thick line – mean; shaded region – standard deviation). (D) The average number of localized point defects of each type (+1, −1/2 and +1/2) at different stages of fragment regeneration (shaded region – standard deviation). The configuration of point defects develops over time. Inset: the net charge of localized point defects as a function of time. The net charge reaches +2 (the topologically required value) only at ~24 hours, after the ordered regions expand and defects become localized. At earlier times (<24 hours), the sum of charge from all the point defects is less than +2 (due to the extended disordered regions). The defect distribution was determined for a population of regenerating fragments that were imaged using a spinning-disk confocal microscope from 4 directions within square FEP tubes at the specified time-points (*T=*0, 2, 6, 24, 48, 72 hr with *N=*16, 25, 56, 44, 43, 41 samples at each time point, respectively; see Methods). The data refers to defects that arise from the disordered regions and their evolution, and does not account for the additional defects associated with tentacle formation.

The folded spheroid with partial nematic order regenerates into a mature *Hydra* within 2-3 days (Fig. 2; Movie 1). This process involves extensive tissue deformations that are primarily driven by internal forces, generated by the actomyosin cytoskeleton, as well as by osmotic forces that cause changes in fluid volume in the internal gastric cavity, leading to cycles of osmotic swelling and ruptures ^31^. During the first ~24 hours after excision, nematic arrays of actin fibers gradually form in the previously disordered regions (Fig. S5), and the fraction of the spheroids’ surface area that contains ordered fiber arrays increases to span nearly the entire surface (Fig. 2B,C). The order appears to “invade” the disordered regions, as the orientation of newly formed fibers is influenced by the alignment of neighboring regions (Fig. S5). As previously shown, this induction of order confers a structural memory of the original body axis orientation ^6^.

Due to the topological constraint of the closed surface, fiber alignment cannot be continuous everywhere and topological defects in the nematic order must emerge (Fig. 2A). As the extended disordered regions progressively become ordered, the topological defects become spatially-localized and develop into well-defined point defects with charges of +1/2, −1/2 or +1 (Figs. 1G-I, 2B,D). After ~24 hours, the total charge of all the point defects in a regenerating spheroid reaches +2 (Fig. 2D inset). While this total charge (following the induction of order) is dictated by topology, the defect configuration that emerges is not. Moreover, the initial defect configuration that appears in the regenerating spheroids is generally different from the final defect configuration in regenerated *Hydra*. For example, the common defect type in regenerating spheroids originating from excised tissue fragments at 24 hours after excision is a +1/2 defect (Fig. 2D), which is entirely absent in mature *Hydra* (Fig. 1). Over time, +1 defects become more prevalent and the +1/2 defects become less frequent, until eventually they are completely eliminated (Fig. 2D). Importantly, the defect configuration in regenerating spheroids depends on the geometry of the excised tissue, because of the influence of the excised geometry on the folding process and the initial actin fiber organization in the folded spheroid ^6^. For example, tissue rings excised from the tubular body of a mature *Hydra* can form a spheroid with two +1 defects from the onset, as the top and bottom faces of the ring seal ^6^, whereas spheroids that originated from excised open rings that fold asymmetrically ^6^ exhibit a much higher incidence of −1/2 defects (Fig. S6B). The different defect configurations that emerge in regenerating tissues originating from different excised geometries evolve and typically stabilize into the stereotypical defect configuration of a mature *Hydra* with +1 defects at the tip of the head and the foot.

## Defect dynamics and the emergence of morphological features in regenerating Hydra tissues

To understand how the defect configuration evolves, we next sought to follow actin fiber dynamics in regenerating *Hydra* and track defect positions over time. This goal is challenging due the extensive tissue movements and deformations that naturally occur during *Hydra* regeneration. To tackle this problem, we developed a method to label specific tissue regions by introducing a cytosolic photoactivatable dye into the ectodermal layer of regenerating tissues by electroporation and using a confocal microscope to locally uncage the dye at well-defined regions (Methods). The photoactivatable dye remains within the cells of the uncaged region, providing a fiducial marker in the regenerating tissue (Fig. S5C). Using this technique, we examined the movement of defects relative to labeled regions throughout the regeneration process. This allowed us to characterize the dynamics displayed by defects of different types, relative to the underlying cells, and relate them to the emergence of morphological features and body plan formation (Figs. 3,4).

**Fig. 3.**
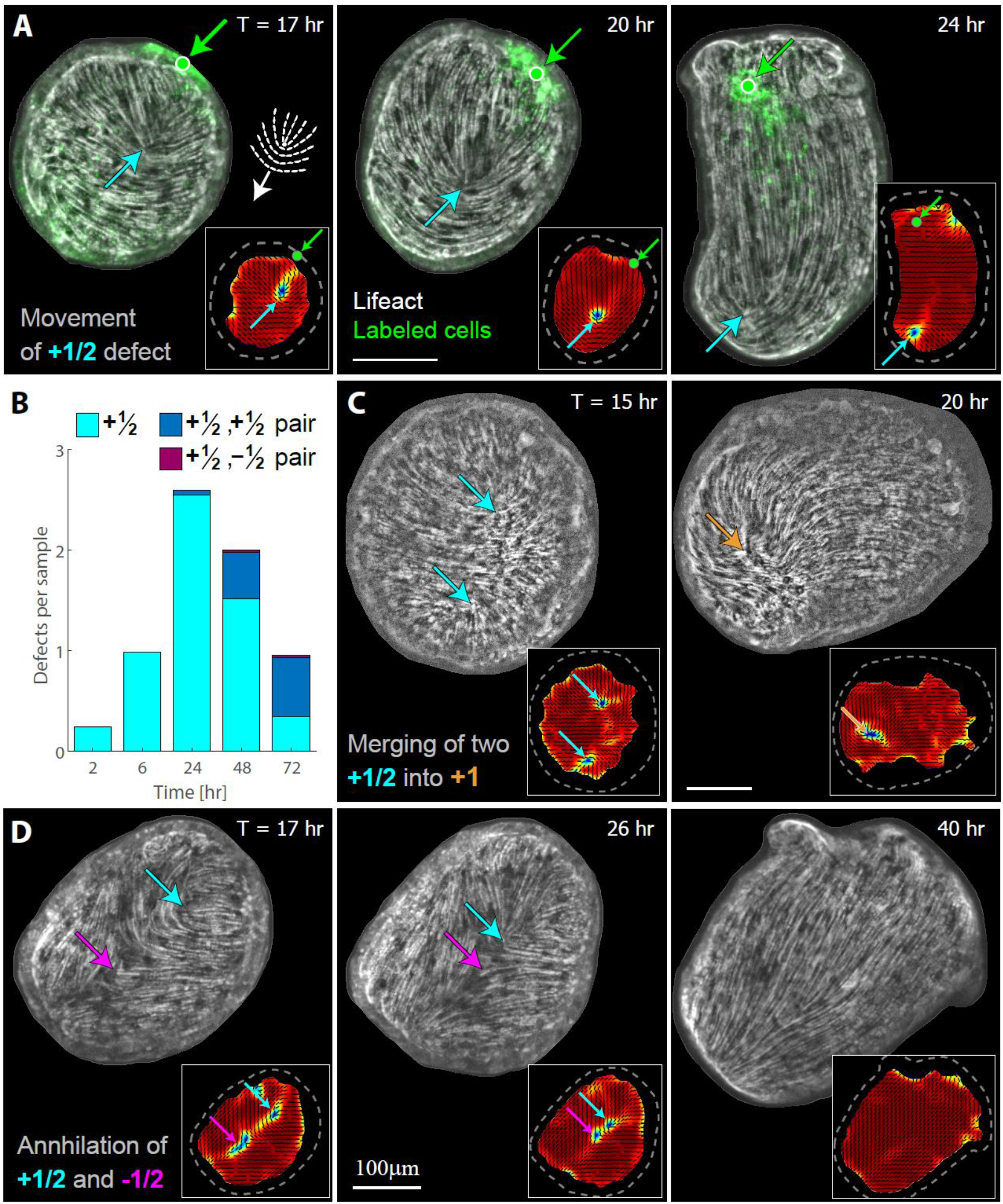
+1/2 defects in regenerating *Hydra* are mobile and can be eliminated. (A) Projected images from a spinning-disk confocal time-lapse movie depicting the movement of a +1/2 defect relative to the underlying tissue (Movie 2). The lifeact-GFP signal (gray scale) is overlaid with the fluorescent tissue label (green). The +1/2 defect (cyan arrow) moves away from the labeled tissue landmark (green arrow) in the direction of its rounded end. (B) Bar plot showing the number of +1/2 defects in regenerating tissue fragments as a function of time. The number of defects which are part of a +1/2, +1/2 pair or a +1/2, −1/2 pair, and hence expected to undergo merging or annihilation, respectively, is indicated. The data was extracted from regenerating fragments that were imaged using a spinning-disk confocal microscope from 4 directions within square FEP tubes at the specified time-points (*T=*2, 6, 24, 48, 72 hr, with *N=* 25, 56, 44, 43, 41 samples at each time point, respectively; see Methods). Two defects were considered a pair when the distance between their cores was smaller than 40 μm. Note that the data refers only to +1/2 defects and hence does not reflect the total charge in the system. (C) Projected images from a spinning-disk confocal time-lapse movie showing the merging of two +1/2 defects into a +1 defect. (D) Projected images from a spinning-disk confocal time-lapse movie showing the annihilation of a +1/2 defect and a −1/2 defect (Movie 3). Insets in (A), (C) and (D) depict maps of the corresponding actin fiber orientation (black lines) and the local order parameter (color). The tissue outlines are indicated with dashed lines.

**Fig. 4.**
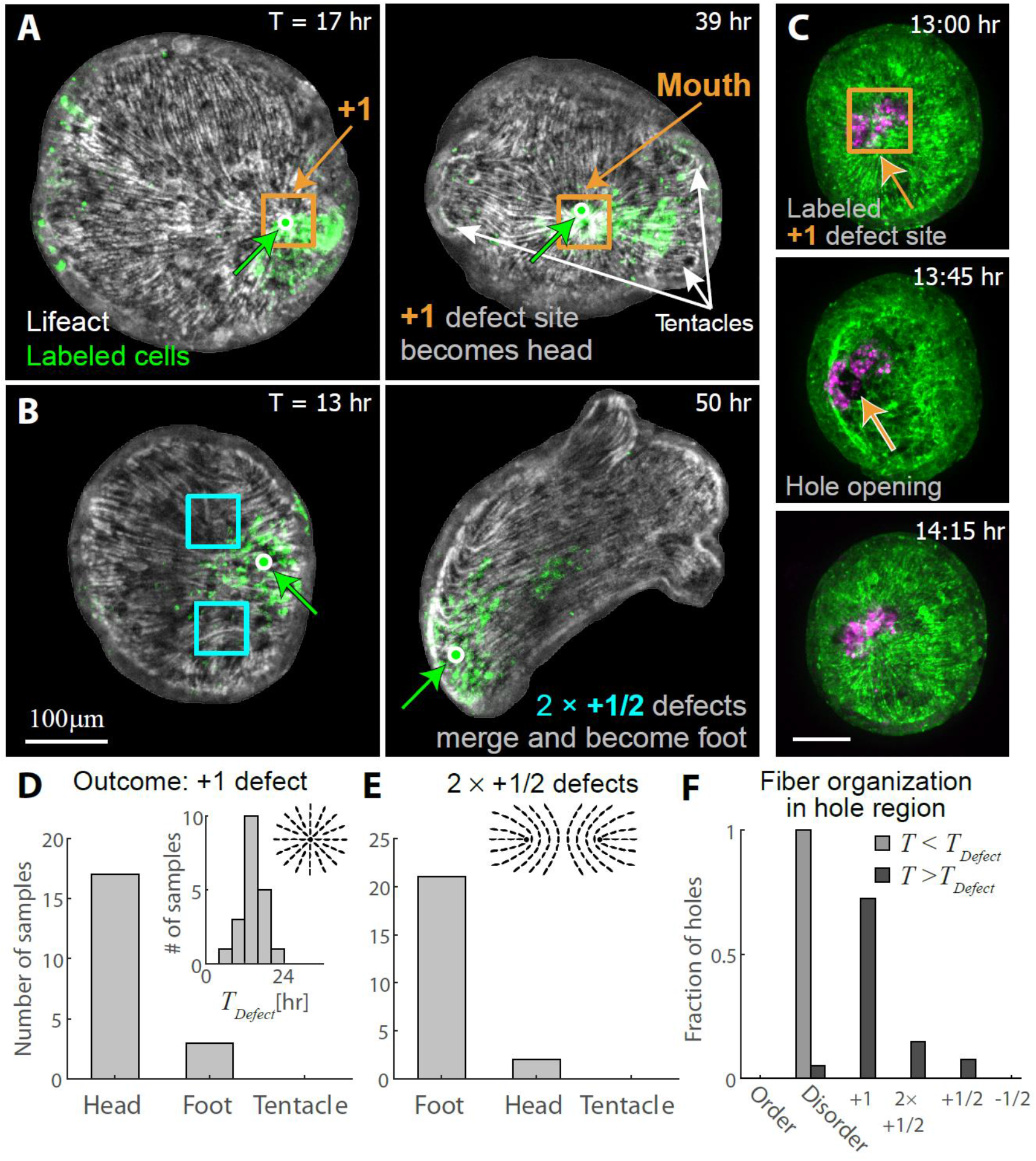
+1 defects as organization centers of head and foot regeneration in *Hydra*. (A) Projected images from a spinning-disk confocal time-lapse movie depicting a +1 defect site that becomes the tip of the head of a regenerated *Hydra* (Movie 4). The lifeact-GFP signal (gray scale) is overlaid with the fluorescent tissue label (green). The +1 defect remains localized adjacent to the labeled tissue throughout the regeneration process, until the region eventually develops into the mouth of the regenerated animal. The position of the defect and the labeled region are indicated. (B) Projected images from a spinning-disk confocal time-lapse movie depicting two +1/2 defects, facing each other (left), that eventually merge to form a +1 defect and become the apex of the foot of the regenerated animal (Movie 5). The lifeact-GFP signal (gray scale) is overlaid with the fluorescent tissue label (green). The position of the defects and a landmark in the labeled region are indicated. (C) Projected images from a spinning-disk confocal time-lapse movie depicting the formation of a rupture hole at a +1 defect site. The image sequence shows the labeled +1 defect site prior (top), during (middle) and after (bottom) hole formation. The lifeact-GFP signal (green) is overlaid with the fluorescent tissue label (magenta). (D,E) The time evolution of defect sites was followed in 44 regenerating *Hydra* samples, labeled by uncaging of a photoactivatable dye or using adhered beads to generate fiduciary landmarks that could be followed over time (Methods). The outcome of the tissue regions surrounding +1 defects that were formed either directly from the disordered region as in (A), or from merging of two +1/2 defects facing each other as in (B), are depicted in (D) and (E), respectively. Typically, since the samples are imaged from one side, we could only study one of the two +1 defect sites in each sample. The inset in (D) shows the distribution of time after excision when localized +1 defects become apparent. (F) The fiber organization surrounding sites of hole formation was determined from time-lapse spinning-disk confocal movies of 27 regenerating tissues. For each sample the time after excision when localized point defects become apparent was determined. The fiber organization surrounding rupture sites was then analyzed for holes that were observed before (*N*=31; light gray) and after the appearance of localized point defects (*N*=39; dark gray), all within the first 24 hours after excision.

We find that +1/2 defects, which are prevalent early in regeneration from tissue fragments (Fig. 2D), are mobile and susceptible to elimination (Fig. 3). We observe +1/2 defects moving relative to the underlying tissue, in the direction of their rounded end (Fig. 3A; Movie 2). These +1/2 defects, that appear during the initial 24 hours after excision, are eliminated over time (Fig. 3B). Their elimination occurs most commonly by merging of two +1/2 defects into a +1 defect (Fig. 3C), or alternatively by annihilation with a −1/2 defect (Fig. 3D; Movie 3). The motility of +1/2 defects, as observed in many other active nematic systems ^19–22, 24–26, 32, 33^ and their two accessible modes of elimination, explain their inherent instability in the tissue and their absence in mature *Hydra*.

In contrast, we find that +1 defects are stable and can be related to the emergence of the head and foot in regenerated animals (Fig. 4). The +1 defects originate either from newly-aligned fibers in the initially disordered regions, or from merging of two previously-formed +1/2 defects (Fig. 3C). Typically, a +1 defect forms in the previously disordered regions within the first 24 hours, sometimes as early as 3-4 hours after excision (Fig. 4D inset). These +1 defects remain stationary relative to the underlying tissue and can be followed throughout the regeneration process, at the end of which they will predominantly be located at the mouth of the regenerated animal (Fig. 4A,D; Movie 4). Alternatively, +1 defects can arise through merging, when two +1/2 defects aligned with their curved ends facing each other, move closer together as the tissue elongates until coalescing, most typically to form the foot (Fig. 4B,E; Movie 5). The final merging often occurs late in the regeneration process, after the formation of tentacles and the establishment of the body axis.

Together, these findings indicate that it is possible to discern the sites where +1 defects will emerge, already at very early stages of the regeneration process. These sites remain stationary with respect to the tissue and hence identify the region of cells that will eventually form the functional morphological features of the regenerated animal (Movies 4, 5). Importantly, there is a clear asymmetry between the head and foot forming processes: early +1 defects become the site of head formation, whereas two +1/2 defects merge at the site that becomes the foot of the regenerated animal (Fig. 4). This asymmetry is far from trivial, and is not an inevitable consequence of the topological constraint on the nematic field. In particular, while a +1 defect and two +1/2 defects are topologically similar (having a net charge of +1), the local structure and dynamics at these sites are different, leading to distinct local mechanical environments. Specifically, cells at the core of a +1 defect are positioned at the core of an aster of contracting actin fibers throughout the regeneration process, whereas the cells located between two +1/2 defects (i.e. at the region that will become the foot, after the two +1/2 defects merge) are positioned within regions containing well-ordered fibers over much of the regeneration process. Remarkably, this asymmetry between the head and foot forming regions with respect to their initial defect configuration (Fig. 4D,E), implies that the sites of emergence of the head and the foot in regenerating fragments and the body axis polarity can be predicted long before the physical appearance of these features, solely based on the distribution of actin fiber alignment shortly after excision.

Although −1/2 defects are sometimes observed early in tissue spheroids, they are relatively rare in samples regenerating from excised square tissue fragments (Fig. 2D), and can be eliminated through annihilation with +1/2 defects (Fig. 3D). Early −1/2 defects that survive can end up defining the junction of an additional body axis in multi-axis animals (Fig. S6). The prevalence of early −1/2 defects depends on the initial excised tissue geometry; In particular, spheroids regenerating from excised open rings exhibit a higher average number of −1/2 defects (Fig. S6B) compared to spheroids regenerating from square tissue fragments (Fig. 2D), in correlation with the higher incidence of multi-axis animals in regeneration from open rings ^6^.

Most commonly, however, −1/2 defects emerge much later in regeneration, after the polar body axis is well-defined, during the formation of tentacles. *De novo* defect formation (also known as defect unbinding) typically involves the coupled appearance of two −1/2 defects and a +1 defect (with net zero charge), in conjunction with an increase in surface curvature and the formation of a “bump” that develops into the tentacle (Fig. S7). This process is analogous to that observed during budding, where two −1/2 defects emerge at the base of the early bud together with a +1 defect at its tip, as the bud begins to protrude ^5^. In both cases, the emergence of defects and the appearance of a local protrusion occur simultaneously (Fig. S7)^5^. These defect formation processes in *Hydra* differ from other recently studied active nematic systems, where new defects typically appear as individual +1/2 and −1/2 defect pairs^19–22, 24, 32–34^. Apart from the protrusion-coupled emergence of defects in the tentacles (Fig. S7), we do not observe spontaneous generation of defects in ordered regions during regeneration. Importantly, this means that the configuration of defects that establishes the regenerating animal’s body plan, and defines its head and foot regions, is the result of the evolution and interaction of defects emerging from the disordered regions early in regeneration following the induction of order there.

Interestingly, we have also found an intriguing relation between the nematic fiber organization, the morphological outcome of the region, and the appearance of transient rupture holes which allow the release of pressure and fluid from the spheroid cavity ^31, 35^. The formation of holes in regenerating spheroids is never observed within regions containing ordered nematic fiber alignment (Fig. 4F). Rather, during the earlier stages of regeneration, before localized point defects develop, the ruptures occur in disordered regions (Fig. 4F). Most commonly, the holes appear in disordered regions that are surrounded by ordered fibers that create an effective, spread out +1 “aster”-shaped defect, and proceed to evolve into localized +1 defects as the regions containing ordered fibers expand (Fig. S8). At later times, after localized defects appear, hole formation occurs almost exclusively at +1 defect sites (Fig. 4C,F).

The localization of the tissue rupture sites at +1 defects indicates that the mechanical environment at these sites is significantly altered relative to the surrounding ordered regions. The unique mechanical environment at +1 defect sites appears inherent to their structure as the focal point of an aster of contractile actin fibers. The aster fiber arrangement at the defect core implies that there is a lack of fibers spanning cell-cell boundaries there, likely weakening local cell adhesion (in contrast to regions with ordered fibers which strongly reinforce cell-cell adhesions ^5^). Moreover, during tissue contractions, we expect a high level of tension at the defect core, as the focal point of pulling forces applied by the aster of actin fibers contracting from all directions. Both of these effects, namely having weaker adhesion and higher tension, are expected (from a purely structural and mechanical point of view) to make +1 defect sites more prone to rupture. Since the same defects that facilitate tissue rupture are also those giving rise to the formation of the mouth, we suggest that the altered mechanical environment at these locations reinforces the morphogenetic processes that lead to the required biological specialization at this location, with +1 defect sites acting as precursors for mouth formation.

## Discussion

Our main finding here is that topological defects in the nematic actin fiber alignment that are identified early in regeneration, act as organization centers of the *Hydra* body plan in the morphogenesis process. The nematic actin fiber orientation stands out as a relevant observable, i.e. a coarse-grained field whose dynamics provide a valuable phenomenological description of the regeneration process. This nematic orientation field contains only a small number of topological defects that evolve over the course of regeneration and change at rates that are slow compared to the underlying molecular and cellular processes (Figs. 2–3). Our work primarily deals with the establishment of the polar body axis with a head on one side and a foot on the opposite side, and its relation to the defect configuration in the folded spheroid. While the topological constraint on the total charge in a closed spheroid implies that there must be nematic topological defects in the system, it does not dictate the type of defects, their dynamics, nor how the defects relate to the formation of morphological features in the regenerated animal. One of the main results of our work is the clear asymmetry between head and foot formation with respect to the underlying topological defect configuration (Fig. 4). As such, the defect configuration that arises early in regeneration allows us to identify and differentiate between the distinct sites of head and foot formation, long before the appearance of any morphological features. The location of these morphological features also marks the polarity of the animal, which in regenerating tissue fragments cannot be trivially inherited from the parent animal due to the initial tissue folding (Fig. 2). This ability to predict the sites of head and foot formation from the nematic field, strongly suggest that the defect distribution and dynamics are coupled to the mechanisms stabilizing the body-axis alignment and polarity, and thus likely play a role in the establishment of the body plan.

The organization of the contractile actin fibers around topological defects shapes the local mechanical environment at these sites, which we believe reinforces the morphogenetic processes that eventually lead to the formation of morphological features there. The most important morphogenetic event in *Hydra* morphogenesis is the formation of a “head organizer” at the tip of the mouth that emits inducing signals for head formation (as well as inhibitory ones preventing formation of additional heads) ^7^. Following excision of a tissue piece from the gastric region of the parent animal, the original head organizer is lost and the regenerating tissue must develop a new organizer. Our results indicate that the mouth of the regenerated animal, and hence the new organizer, appear at a +1 defect site (Fig. 4D). Moreover, we show that the +1 defects emerge early in regeneration (Fig. 4D inset) and once formed remain stationary with respect to the underlying tissue (Fig. 4A). Thus, the unique mechanical environment at the core of a +1 defect (which arises from its structure as the focal point of an aster of contractile actin fibers) persists for the duration of the regeneration process, providing localized mechanical cues to the same group of cells that differentiate to form the new organizer. While the details of how a new organizer forms in regenerating tissue pieces are still unclear ^7^, we believe that the mechanical environment at the +1 defect site provides important input that feeds into the auto-catalytic signaling network associated with the emergence of a new head organizer ^7^ and promotes the formation of a head specifically at the defect site.

The coincidence between defect locations and the emergence of morphological features with specialized biological functions, demonstrates the innate coupling between biochemical and mechanical fields in the tissue, rather than implying a direct causal sequence of events driven by the presence of defects. For example, early +1 defects are likely important for head formation, even though the simplistic association of the presence of a +1 defect with the formation of a head organizer is obviously wrong (since +1 defects can be present elsewhere e.g. at the tip of the tentacles). Similarly, tentacle or bud formation are not initiated by preexisting defects, but rather involve *de novo* defect formation, likely triggered by biochemical signals ^5^. Nevertheless, the interaction between the mechanical field and biochemical signaling pathways provides a possibility for reinforcement of specific processes at defect sites, with their unique mechanical environment, towards the establishment of the required biological functions there. Moreover, beyond their local effects, topological defects in nematic systems have long-range interactions and are coupled to the geometry of the surface, in addition to the topological constraint on the total defect charge ^16^. As such, topological defects naturally relate global characteristics of the animal’s emerging body plan with local properties of the regions in which morphological features develop, and can thus contribute to the large-scale coordination and robustness of the morphogenesis process.

Based on our results, we suggest a complementary view of morphogenesis, in which a physical field (here, the nematic actin fiber orientation) can be considered as a “mechanical morphogen” field that provides relevant cues for directing the morphogenesis process. This view extends Turing’s framework for the chemical basis of morphogenesis that has been extremely influential in developmental biology ^8, 9^, and suggests a physical field as a mechanical counterpart to the chemical morphogen field. The physical field reflects an integration of multiple non-trivial biochemical and biophysical factors including the mechanical properties, geometry and stress in the tissue, and the complex mechanosensitive response to these features within the tissue. We hypothesize that the interplay between various biochemical signaling processes and this mechanical morphogen field with its associated pattern of topological defects, leads to the establishment of a robust body plan in regenerated *Hydra*. We expect extensive mutual feedback between the mechanical morphogen field and the signaling processes: the nematic fiber organization has a major influence on the stress distribution and shape of the tissue which will influence the distribution of chemical morphogens, whereas signaling processes can modulate actin-myosin dynamics leading to changes in the nematic fiber organization. In the presence of such feedback, it is unlikely that the mechanical or the chemical morphogen fields direct major morphogenetic events on their own. Nevertheless, the identification of defect sites as precursors for morphological features, draws attention to these regions as focal points of the patterning process, offering the opportunity for focused investigation of the synergistic interplay between biochemical signaling pathways and mechanical cues at these locations. The development of an integrated framework for morphogenesis, which incorporates the dynamic interplay between self-organized physical fields and signaling processes, and extends it to other tissues and developing organisms, presents an exciting challenge for future research.

## Acknowledgements

We thank Gidi Ben Yoseph for superb technical assistance. We thank Nitsan Dahan from the LS&E Imaging and Microscopy Unit for help with confocal and light-sheet microscopy. We thank Prof. Bert Hobmayer for generously providing transgenic Hydra expressing lifeact-GFP. We thank Vincenzo Vitelli, Christina Marchetti and Julia Yeomans for valuable discussions. We thank Natalie Dye and Meghan Driscoll for advice on image analysis. We thank Pascal Silberzan, Alex Mogilner, Jacque Prost, Natalie Dye, Yariv Kafri, Tom Schultheiss, Guy Bunin, Anna Frishman and Niv Ierushalmi for comments on the manuscript.

This work was supported by a grant from the European Research Council (ERC-2018-COG grant 819174) to K.K., a grant from the Israel Science Foundation (grant No. 228/17) to E.B., and a Miriam and Aaron Gutwirth Memorial Fellowship to Y.M.S.

## Methods

### Hydra strains, culture and sample preparation

All the experiments are performed using transgenic *Hydra* strains expressing lifeact-GFP in the ectodermal cells, generously provided by Prof. Bert Hobmayer from the University of Innsbruck, Austria. Animals are cultured in standard *Hydra* culture medium (HM) (1mM NaHCO3, 1mM CaCl2, 0.1mM MgCl2, 0.1mM KCl, 1mM Tris-HCl pH 7.7) at 18° C. The animals are fed 3-4 times a week with live Artemia Nauplii and rinsed after ~4 hours. Tissue segments are excised from the middle body section of mature *Hydra*, ~ 24 hours after feeding, using a scalpel equipped with a #15 blade. Tissue rings are excised by performing two nearby transverse cuts. To obtain tissue fragments, a ring is cut into ~4 parts by additional longitudinal cuts.

### Sample Preparation

To reduce tissue movements and rotations, samples are either embedded in a soft gel (0.5% low melting point agarose (Sigma) prepared in HM), or placed in HM supplemented with 1% Methyl Cellulose. The general characteristics of the regeneration process in these environments are similar to regeneration in normal aqueous media. For experiments in gels, the regenerating tissues are placed in liquefied gel ~2-6 hours after excision (to allow the tissue pieces to first fold into spheroids), and the gel subsequently solidifies around the tissue.

Time-lapse spinning-disk confocal imaging is done on samples placed in 35mm glass-bottom petri dishes (Fluorodish), or polymer coverslip 24-well plates (Ibidi μ-plate 24 well, tissue culture treated), embedded in 2-3 mm of gel that is layered with ~3-4 mm of HM from above. Light-sheet microscopy samples are loaded in liquefied gel into a ~1cm long cylindrical FEP tube with an internal diameter of 2.15 mm (Zeiss Z1 sample preparation kit). Once the gel has set, the tube is mounted on one end onto a glass capillary which is held by the microscope sample holder. The imaging is done through the FEP tubing, which has a similar refractive index to the surrounding solution. Movies from 4 different angles are acquired by rotating the specimen.

To image a large number of samples from all directions by spinning-disk confocal microscopy, regenerating tissues are similarly loaded into FEP tubes with a square cross-section and an internal diameter of 2 mm (Adtech). The tubes are rotated manually to each of the 4 flat facets of the square tube and secured using a homemade Teflon holder to keep them stationary at each orientation. Images from 4 directions are acquired at the specified time points.

### Microscopy

Spinning-disk confocal z-stacks are acquired on a spinning-disk confocal microscope (Intelligent Imaging Innovations) running Slidebook software. The lifeact-GFP is excited using a 50mW 488nm laser and the activated Abberior CAGE 552 is excited using a 50mW 561nm laser. Imaging is done using appropriate emission filters at room temperature, and acquired with an EM-CCD (QuantEM; Photometrix). Long time lapse movies of regenerating Hydra are taken using a 10× air objective (NA=0.5). Short time lapse movies and images in FEP tubes are taken using a 10× water objective (NA=0.45).

Light-sheet microscopy is done on a Lightsheet Z.1 microscope (Zeiss). The light-sheet is generated by a pair of 10× air objectives (NA=0.2), imaged through 20× water objectives (NA=1), and acquired using a pair of CMOS cameras (PCO.edge). The lifeact-GFP is excited using a 50mW 488nm laser and the activated Abberior CAGE 552 is excited using a 50mW 561nm laser. The imaging is performed using the “pivot scan” setting to minimize imaging artefacts that introduce streaking in the direction of illumination, yet some remnants of the streaking artefacts are still apparent in the images.

### Tissue labeling using photoactivation of caged dyes

To label specific tissue regions we use laser photoactivation of a caged dye (Abberior CAGE 552 NHS ester) that is electroporated uniformly into mature *Hydra* and subsequently activated in the desired region. Electroporation of the probe into live *Hydra* is performed using a homemade electroporation setup. The electroporation chamber consists of a small Teflon well, with 2 perpendicular Platinum electrodes, spaced 2.5 mm apart, on both sides of the well. A single *Hydra* is placed in the chamber in 10μl of HM supplemented with 2mM of the caged dye. A 75 Volts electric pulse is applied for 35ms. The animal is then washed in HM and allowed to recover for several hours to 1 day prior to tissue excision.

Following excision, the specific region of interest is activated by a UV laser in a LSM 710 laser scanning confocal microscope (Zeiss), using a 20× air objective (NA=0.8). The samples are first briefly imaged with a 30 mW 488 nm multiline argon laser at 0.4-0.8% power to visualize the lifeact-GFP signal and identify the region of interest for activation. Photoactivation of the caged dye is done using a 10 mW 405nm laser at 100 %. The activation of the Abberior CAGE 552 is verified by imaging with a 10 mW 543 nm laser at 1%. Subsequent imaging of the lifeact-GFP signal and the uncaged cytosolic label is done by spinning-disk confocal microscopy or light-sheet microscopy as described above. The uncaged dye remains within the cells of the photoactivated region and does not get advected or diffuse away within our experimental time window (Fig. S5C).

### Experiments on hole formation

To follow the formation of holes in regenerating *Hydra* tissue and visualize the expulsion of fluid from the holes, we incubate tissue segments immediately after excision in a solution of fluorescent beads (0.2 μm Fluorospheres, Carboxylate-modified Microspheres, dark red 660/680, Invitrogen), so that as the tissue folds and seals into a hollow spheroid, the internal fluid will contain beads. The beads are first blocked by incubation in a 3% solution of Bovine Serum Albumin (Sigma) in HM, whilst shaken for one hour, and then sonicated for 2 minutes. The bead solution is then further diluted 1:10 in HM. Tissue segments are added to the bead solution immediately after excision and incubated for 3 hours to allow tissue folding and sealing. Tissue spheroids are then rinsed 3-6 times in HM, and imaged in wells (diameter 1 mm) prepared from 2% Agarose gel (Sigma). Imaging was done in a solution of HM containing 1% Methyl Cellulose (Sigma) in order to increase the viscosity of the solution and allow more precise visualization of tissue and bead movement. Some of the beads adhere to the tissue (especially around the closure region), providing fiduciary landmarks on the regenerating tissue.

### Image analysis

The image analysis tools used for extracting the 2D surface geometry of the tissue, the nematic director field and the localization of the activated tissue label are based on custom-written code, as well as adaptation and application of existing algorithms, written in Matlab and ImageJ as detailed below.

#### Layer analysis

The regenerating *Hydra* tissue spheroids consist of a bilayered epithelium surrounding a hollow cavity. The 2D apical and basal surfaces of the ectoderm are computationally identified in the 3D fluorescence images of the lifeact-GFP signal in regenerating tissue spheroids. The supra-cellular actin fibers reside on the basal surface of the ectoderm while the cortices marking the cell outlines are visible on the apical surface. 2D projected images of the basal and apical surfaces of the ectoderm are automatically extracted from the 3D spinning-disk z-stack acquired with a z-interval of 3-5 μm using the “Minimum Cost Z surface Projection” plugin in ImageJ (https://imagej.net/Minimum_Cost_Z_surface_Projection). The cost images are generated by processing the original z-stacks using custom-code written in Matlab. First, the signal from the ectoderm layer was manipulated to make it more homogenous within the layer without increasing its thickness, by applying the built-in Matlab anisotropic diffusion filter.

Subsequently, we employ a gradient filter to highlight the apical and basal surfaces as the top and bottom boundaries of the ectoderm layer. The apical and basal surfaces are determined using the minCost algorithm (Parameters used: Rescale xy: 0.5, rescale z: 1, min distance: 15μm, max distance: 45μm, max delta z: 1, two surfaces). The surfaces given by the minCost algorithm are then smoothed by applying an isotropic 3D Gaussian filter of width 1 pixel (after rescaling to an isotropic grid matching the resolution in the z-direction) and selecting the iso-surface with value 0.5.

#### Image Projection

2D projected images of the ectodermal actin fibers (that reside on the basal surface of the ectodermal layer, Fig. S1) in spinning-disk confocal images are generated by extracting the relevant fluorescence signal from the 3D image stacks based on the smoothed basal surface determined above. To obtain the projected 2D image value for each x-y position, we employ a Gaussian weight function in the z-direction with a sigma of 3 μm, which is centered at the z-value determined from the smoothed basal surface with a small (2-3 pixels) fixed offset found to optimally define the desired surface. 2D projected images of the ectodermal actin fibers in light-sheet movies are generated by taking a maximum intensity projection of the 3D image stacks, which result in better or similar quality projection images compared to method described for spinning-disk images. The resulting 2D projected images are further subject to contrast limited adaptive histogram equalization (CLAHE) in Matlab as described below.

2D projection of spinning-disk confocal images of the photoactivated tissue label are generated by taking a maximum intensity projection of the 3D image stacks in the z-region between the smoothed apical and basal surface determined above.

#### Fiber orientation analysis

A system with nematic orientation is a system in which the constituents tend to align parallel to each other, with no distinction between opposite ends (as opposed to a polar system). The average orientation of the constituents can be described by a director field, which is oriented along the mean orientation determined in a small region surrounding every point in space. Here, the nematic director field describes the local orientation of the supra-cellular ectodermal actin fibers. The nematic director field is extracted in an automated manner from the 2D projected images of the actin fibers using an algorithm that is based on analysis of the local image intensity gradients. Within an image of an ordered fiber array, the changes in image intensity will be strongest perpendicular to the fibers, whereas the intensity along the fibers remains more uniform. The local direction of the image intensity gradient will thus tend to be oriented perpendicular to the fibers, and can hence be used to determine the local fiber orientation. Similar techniques are used, e.g. to identify the ridge orientation in finger print images ^36^.

The local fiber orientation is determined from the smoothed signal intensity gradients in the image as described in ^36^ and implemented in Matlab according to “FingerPrint Matching: A Simple Approach” toolbox by Vahid K. Alilou from the Matlab File Exchange Central (https://www.mathworks.com/matlabcentral/fileexchange/44369-fingerprint-matching-a-simple-approach) with small modifications. All the relevant length scales below are given in pixels, assuming a calibration of 1.28 μm/pixel (as in our typical spinning-disk images). For images with different calibration, the number of pixels is adjusted to maintain the same distances in real spatial units

The signal intensities in the images are first normalized employing CLAHE with Matlab’s “adapthisteq” function with Rayleigh distribution and a tile size of 20 pixels. The signal intensity gradients 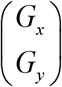 in the normalized images are calculated by convolving the adjusted image, *G*(x,y), with the gradient of a Gaussian with a width of 0.5 pixel. The covariance matrix of the gradient is then calculated from the signal intensity gradient (after averaging using a Gaussian filter with σ=5 pixels), as:

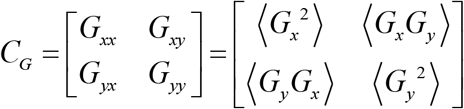

The raw orientation angle is calculated from the covariance matrix as:

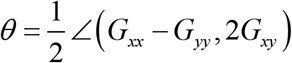

Where ∠(*x*, *y*) is defined as:

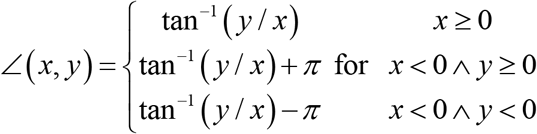

The coherence field, used to determine the extent of nematic order, is calculated from the raw orientation field according to: 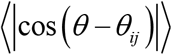, with *θ*_*ij*_ the orientation of a given pixel, and the average is taken over angles within a square window of 20 pixels. The coherence field thus provides a measure of the local variation in the orientation field. Regions with an ordered fiber array have a well-defined orientation locally, so this variation is small and the coherence will be close to 1 (since 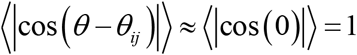), whereas in disordered regions this variation will be higher and the coherence values will be lower. The orientation field is further smoothed by applying a Gaussian filter (σ=3 pixels) on the raw orientation field.

The orientation and coherence fields are defined within a masked area that excludes the regions close to the tissue edges, as the sharp signal gradients near the edges introduce artefacts into the orientation field.

The nematic local order parameter, which is a measure of the local orientational order, is defined as:

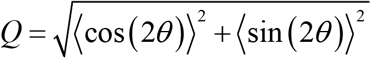

Where *θ* is the orientation field and the averaging is done over a window size of 32 pixels. The nematic order parameter is used to identify the sites of topological defects. In regions which have well-ordered fiber array, the nematic order parameter will be close to 1 (i.e. if *θ*≈*θ*_0_ within a region of size 32 pixels (or ~40μm) then 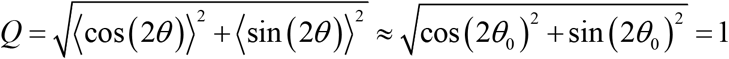). In the vicinity of defects, the nematic orientation varies which will translate to a local minimum in the nematic order parameter at the defect site.

#### Defining regions with ordered fibers

To automatically identify regions containing ordered actin fibers in our images, we define a threshold value for the coherence field (defined in the previous section), above which the region is considered ordered. The coherence threshold (taken as 0.95) is used to create a binary image. The thresholded region in the binary image is subsequently simplified by smoothing with a kernel of size 32 pixels and selection of the contour with value 0.5. The fraction of area (after smoothing) out of the total area of the coherence field is taken as the fraction of the tissue surface with ordered fibers (Fig. 2C).

#### Definition and identification of topological defects in the fiber orientation field

A nematic system can contain singularities in the orientation field, which are points at which the alignment is locally disordered. These singularities are called nematic topological defects. These topological defects form specific patterns around the defect, and can be characterized by their “topological charge”, or winding number, which is the number of times the director orientation rotates around the center of the defect. For example, the orientation field surrounding a defect with topological charge of +1/2 rotates exactly half a rotation (180 degrees) around the defect center. In a nematic system, the topological charge can be any integer multiple of +1/2, because a rotation of 180 degrees brings the director back to its original orientation. A positive sign refers to counter-clockwise rotation, whereas a negative sign refers to clockwise rotation.

The local order parameter *Q* (see definition above) is a measure of how much the local orientation in a certain region varies. At the center of a topological defect, the local order parameter will receive a minimal value, since the orientation field surrounding this point contains all possible orientations.

In the *Hydra* tissue, we identify topological defects as points around which there is well-defined alignment of fibers, and the orientation rotates around the center of the defect, forming the specific patterns characteristic of nematic topological defects. The identification is based on the calculated fiber orientation field, and the associated local order parameter (*Q*; see definition above). Defects are first manually identified by specifying a rectangular region around the defects in every time frame, using the Matlab Ground Truth Labeler application. The defects are then precisely localized by taking the position of the minima in the local order parameter within the specified rectangular regions and tracked over time.

Our definition of defect type is based on manual inspection, looking both at the fiber organization and at the calculated order parameter (defined by averaging over a length scale of 32 pixels). For +1/2 and −1/2 defects, the defects typically appear as a well-defined single minimum in the local order parameter (calculated over a region of size 32 pixels~40μm). The +1 defects often exhibit a larger core size, and may appear as tandem minima which seem as two adjacent +1/2 defects in the local order parameter field. We define a +1 defect to be a region with an overall charge of +1 within a diameter of 40μm, that does not contain an ordered fiber array within the core region. This stands in contrast to a pair of +1/2 defects which are separated by an ordered region and will be hence considered two separate +1/2 //defects.

#### Analysis of hole formation

Analysis of hole formation is performed manually by identifying frames within time-lapse movies in which a hole opens in the tissue. The criteria for defining a hole include a transient, visible gap in fluorescence in both the lifeact-GFP signal and additional fluorescent signal if exists, as well as change in tissue volume before and after the hole appearance and/or visible expulsion of inserted beads or other material from the hole. Times and locations of observed hole opening events are recorded, and hole opening sites are followed over time (using fiduciary landmarks) to determine the fiber organization and morphological outcomes at these sites.

#### Analysis of +1 defects and outcome

Analysis of +1 defects and outcome is performed manually by identifying and following defects over time in fluorescent time-lapse movies of lifeact-GFP in regenerating *Hydra* with local fluorescent marking using photoactivated dyes (see above). The defects included in the analysis are those that can be reliably tracked (with the aid of the introduced landmarks inserted in the tissue) throughout the regeneration process from their formation until the final morphological outcome. The time defined for defect appearance is the time at which ordered fibers are completely visible in the full region encircling the defect.

## Supplementary Figures

**Figure S1.**
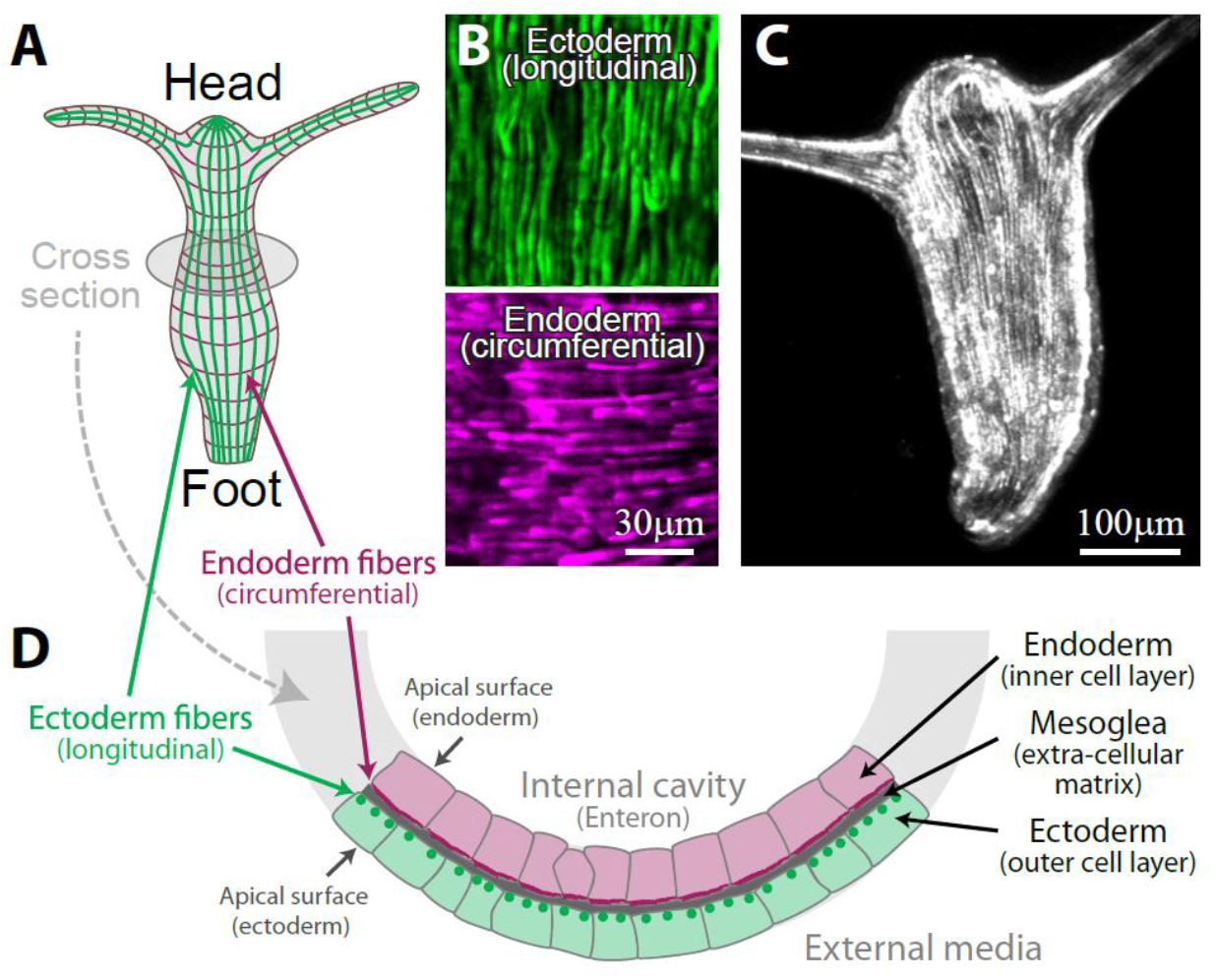
The structure of a mature *Hydra*. (A) Schematic illustration of a mature *Hydra* showing the organization of the supra-cellular actin fibers in the outer ectoderm layer (green) and in the inner endoderm layer (purple). (B) Images of the ectodermal (top) and endodermal (bottom) supra-cellular actin fibers in the body of transgenic *Hydra* expressing lifeact-GFP in the ectoderm or the endoderm, respectively. The fibers in the ectoderm are aligned along the animal axis, whereas the fibers in the endoderm are aligned in a perpendicular, circumferential orientation. We focus in this work on the more prominent ectodermal fibers, which are thicker and appear continuous over supra-cellular scales. (C) Image showing the supra-cellular ectodermal actin fiber organization in a small mature *Hydra* expressing lifeact-GFP. (D) Schematic illustration of a perpendicular cross-section of the tubular *Hydra* body. Part of the ring cross-section is shown, depicting the external ectoderm cell layer, the internal endoderm cell layer, and the extra-cellular matrix (mesoglea) sandwiched between the two layers. The cells in each layer form a polarized epithelial sheet, with their basal surfaces facing the mesoglea, and their apical surfaces facing either the external medium in the ectoderm, or the internal gastric cavity in the endoderm. The ectodermal and endodermal arrays of supra-cellular fibers lie along the basal layer of each epithelial sheet, on a pair of nearly parallel 2D curved surfaces.

**Figure S2.**
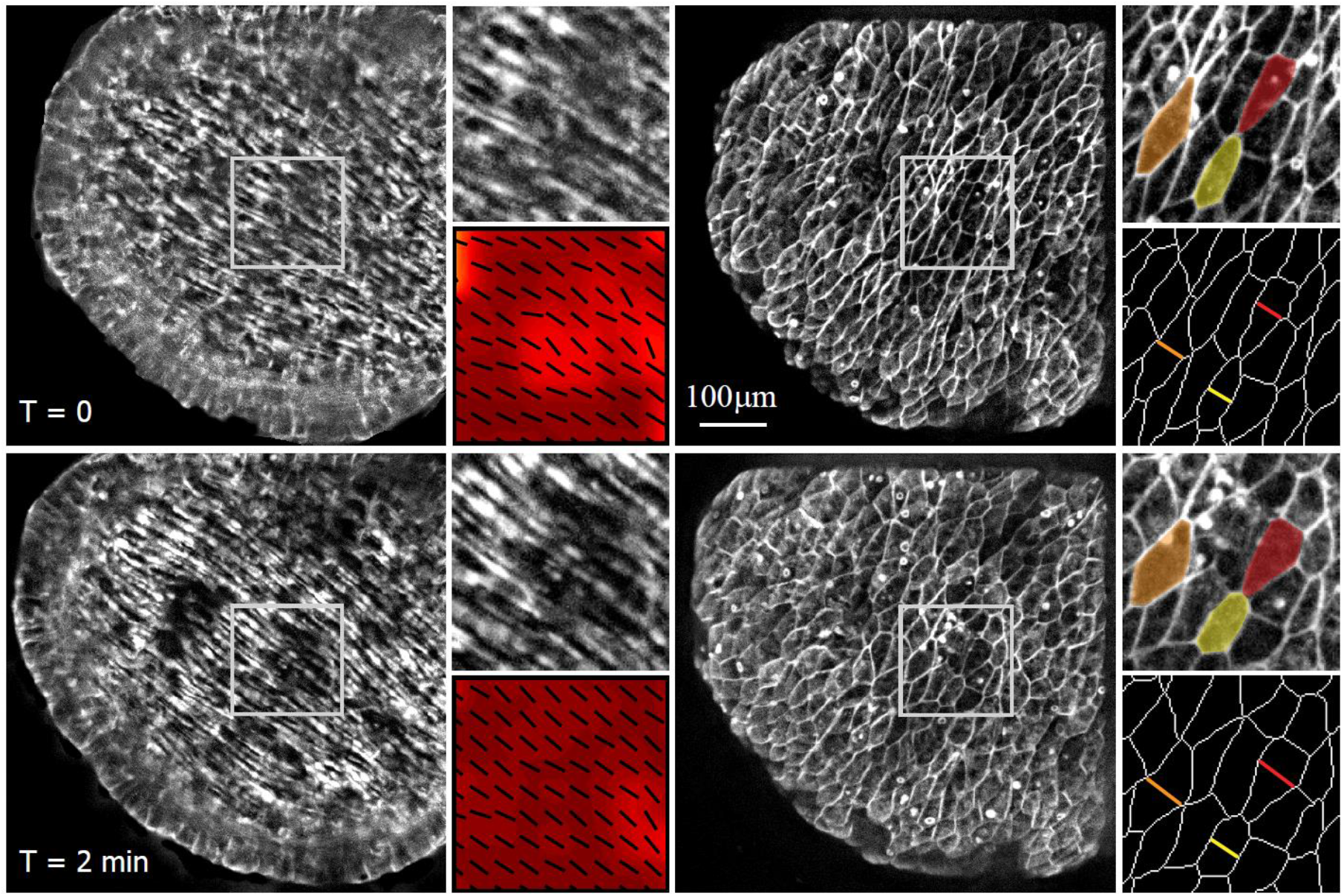
The director field describing the actin fibers in *Hydra* is a slowly-varying in space and time. Images from a time lapse movie of a regenerating tissue fragment expressing lifeact-GFP showing the actin fiber organization at the basal surface of the ectoderm (left) together with the cortical actin at the apical surface of the ectoderm (right). The associated director field (left) and cell outlines (right) extracted from the corresponding zoomed images are also shown (bottom right panels). Three labeled cells are highlighted to illustrate the changes in cell shape over this time interval (2 min). The director field describing the actin fiber orientation varies slowly in space, with typical radii of curvature which are of the order of the tissue size and much larger than the scale of individual cells. Furthermore, while the tissue morphology and cellular shapes change significantly between the two images separated by a time interval of 2 minutes, the nematic orientation field remains stable and changes over much longer time scales (hours).

**Figure S3.**
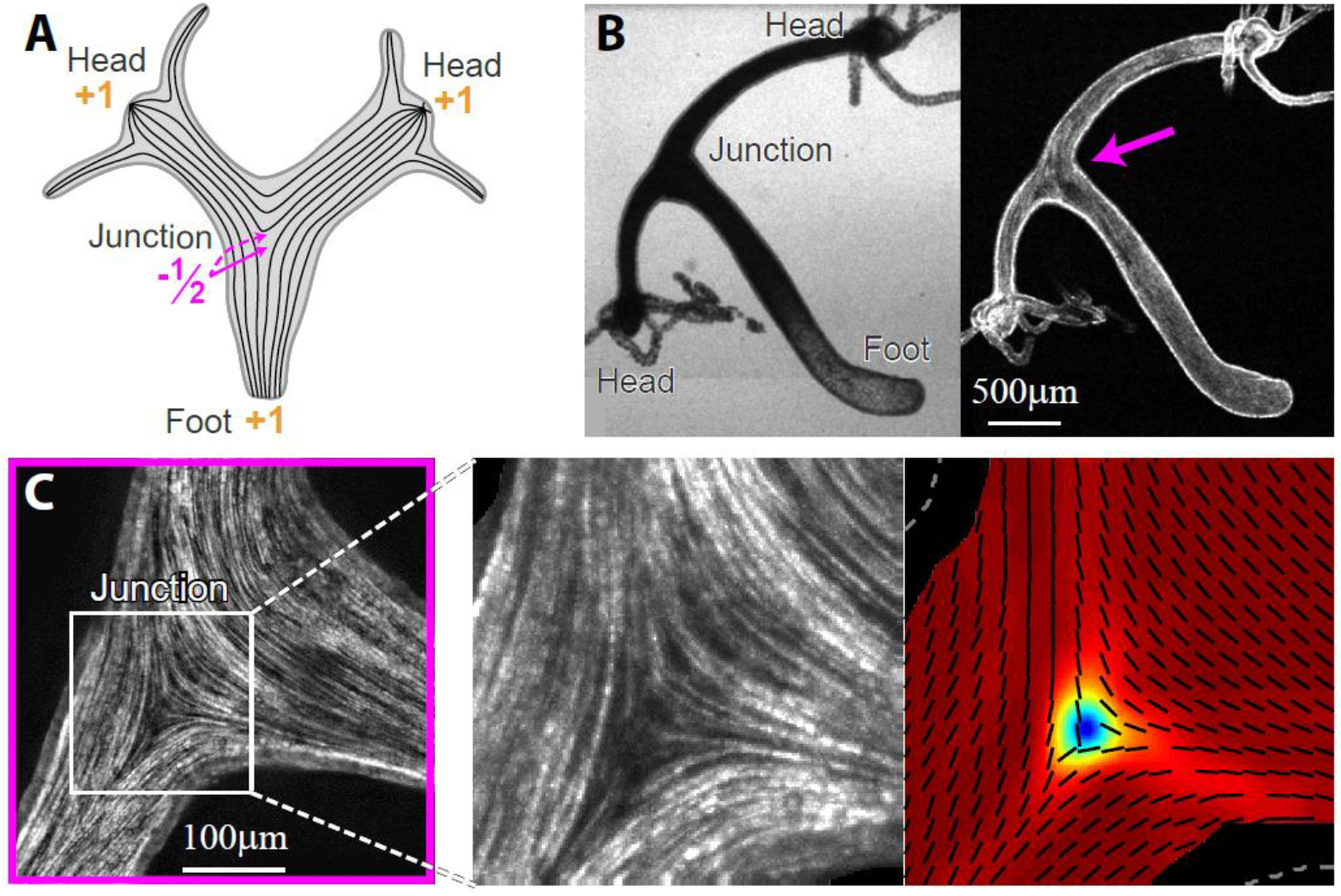
Topological defects in multi-axis *Hydra*. (A) Schematic illustration of the configuration of topological defects in a multi-axis *Hydra* that has two heads and a foot. The two heads and the foot each have a +1 defect at their tip, whereas the junction between the two axes has two −1/2 defects (one visible; the second −1/2 defect is on the opposite side). (B,C) Images of a two-headed transgenic *Hydra* expressing lifeact-GFP in the ectoderm. (B) Bright-field (left) and spinning-disk confocal (right) images showing the entire *Hydra* body with two heads and a foot. The arrow indicates the junction between the two body axes. (C) Zoomed images showing the actin fiber organization near the junction. Left: Projected spinning-disk confocal images showing the actin fibers. Right: Images of the actin fiber orientation extracted from the image intensity gradients (black lines) and the local order parameter (color; see Methods). The −1/2 defect at the junction is clearly evident.

**Figure S4.**
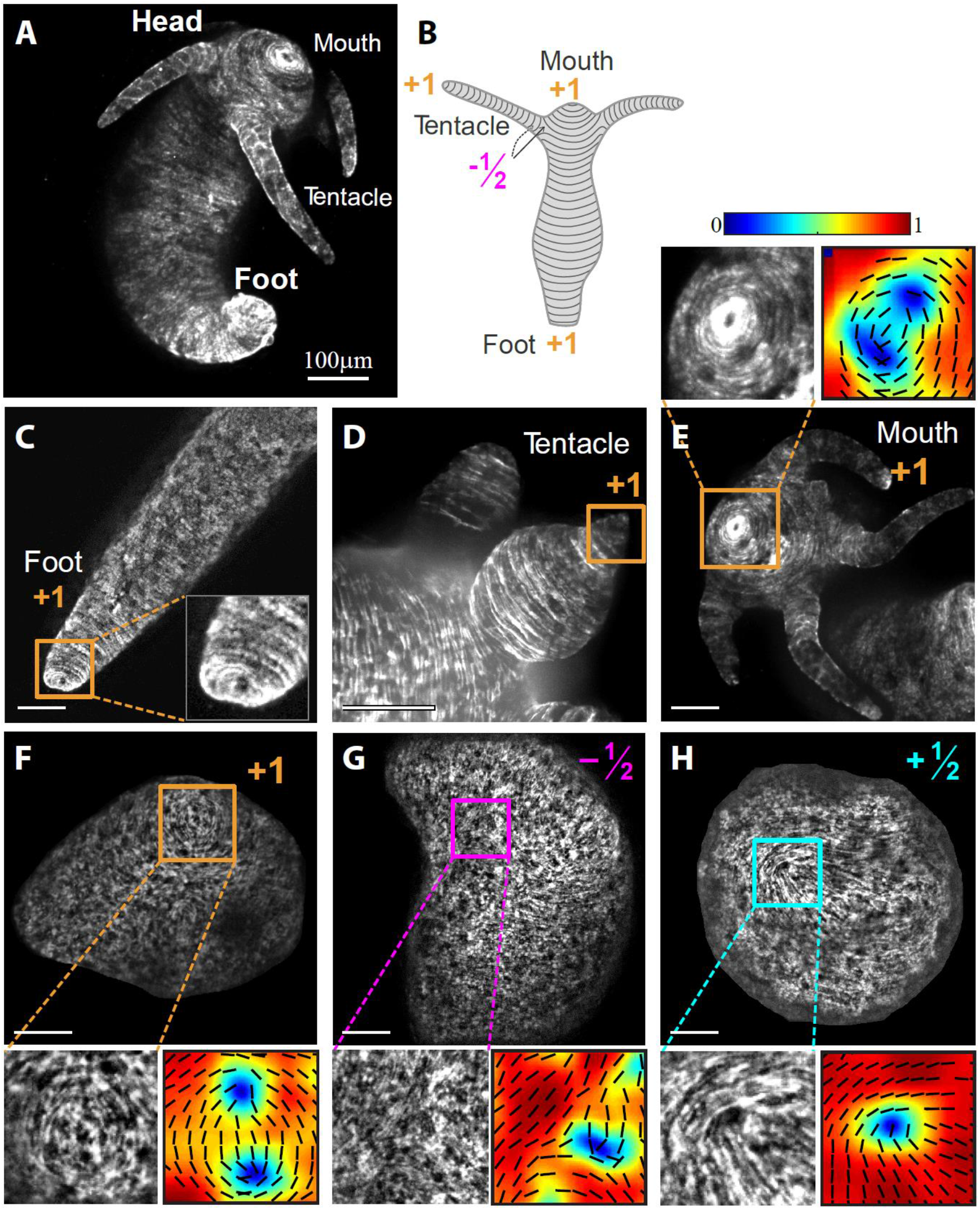
Topological defects in the endoderm layer in *Hydra*. (A) Image showing the nematic actin fiber organization in the endoderm of a small mature *Hydra*. The fibers are oriented in a circumferential orientation along the body axis of the animal, perpendicular to the ectodermal fibers shown in the main text (Fig. 1). (B) Schematic illustration of the nematic actin fiber organization in the endoderm of a mature *Hydra*. The topological defects in the nematic organization of the endodermal fibers coincide with the defects in the ectodermal fibers, localized at the mouth (+1), foot (+1) and tentacles (+1 at the tip, and two −1/2 at the base). (C-E) Images of the endodermal actin fiber organization containing topological defects localized at the functional morphological features of a mature *Hydra*: the base of the foot (C), the tip of a tentacle (D), and the tip of the mouth (E). A zoomed image (top left) and map (top right) in (E) show the vortex-like arrangement of actin fibers around the mouth. The map depicts the fiber orientations extracted from the image intensity gradients (black lines) and the local order parameter (color; see Methods). (F-H) Images showing the different types of topological defects found in the endoderm of regenerating *Hydra* tissue spheroids: +1 defect (F), −1/2 defect (G) and +1/2 defect (H). Insets: zoomed images (left) and maps (right) of the corresponding actin fiber orientation and the local order parameter. All images shown are 2D projections extracted from 3D spinning-disk confocal z-stacks of transgenic *Hydra* expressing lifeact-GFP in the endoderm.

**Figure S5.**
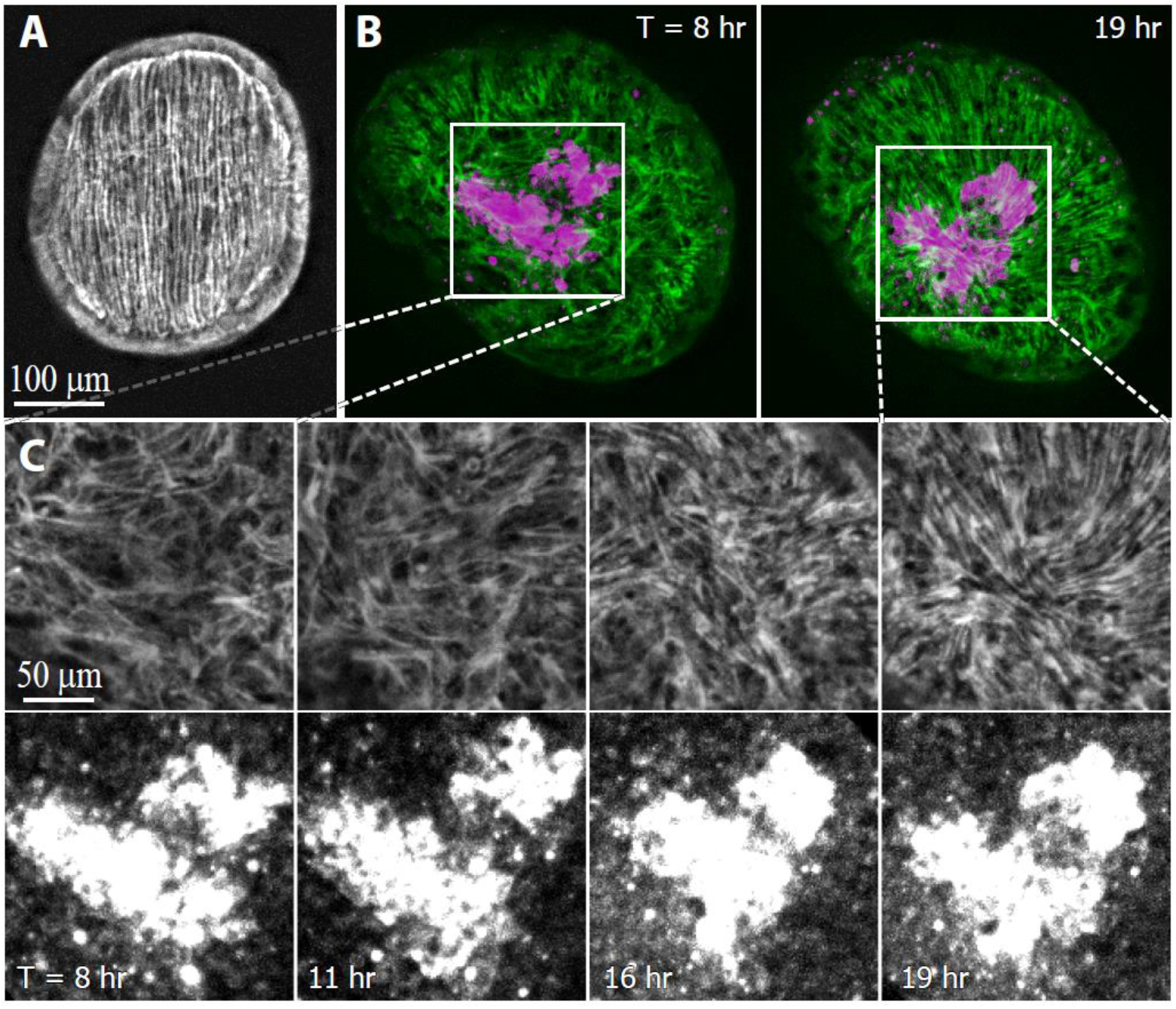
Induction of nematic order in a regenerating tissue. (A) Image of the actin fiber organization in a tissue fragment immediately after excision. The excised fragment is characterized by an open boundary and a complete nematic array of actin fibers inherited from the parent animal. (B) Images from a time-lapse movie of a regenerating tissue spheroid that contains a region labeled by uncaging a photoactivatable dye (Methods). The photoactivated dye is retained within the cells in the uncaged region, allowing us to track this region over time. The ectodermal actin organization (green) and photoactivated tissue label (magenta) are shown at two time points from the movie. (C) Zoomed images showing the actin organization (top) and the photoactivated dye (bottom) around the uncaged region at different time points. Initially, the actin in the folded tissue spheroid exhibits partial nematic order, with a large disordered region around the uncaged region (left). Over time, parallel actin fibers emerge in the disordered region, with an orientation that is aligned with existing fibers in the surrounding tissue. Since the fiber orientation of the surrounding regions is not uniform, localized topological defects appear in the previously disordered region (right). The image sequence depicts steps in this process, as the labeled tissue region transitions from a disordered state to an ordered one, disrupted by localized point defects. All images shown are 2D projections extracted from 3D spinning-disk confocal z-stacks of transgenic *Hydra* expressing lifeact-GFP in the ectoderm.

**Figure S6.**
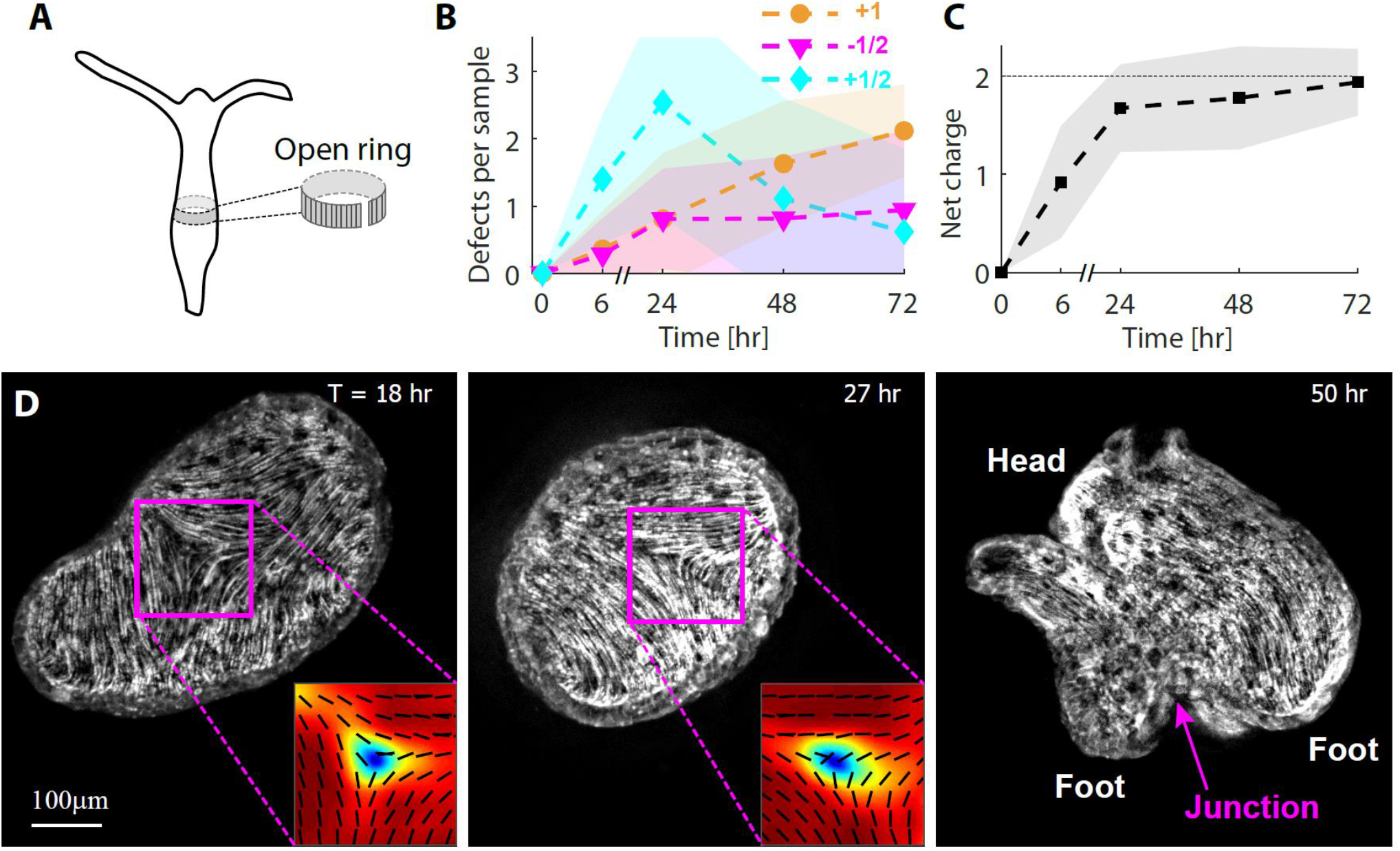
Nematic actin organization in regeneration from excised open rings. (A) Schematic illustration of an excised open tissue ring. The excised tissue ring (which contains an ordered array of fibers) is cut open along the original body axis. We have previously shown that excised open rings fold in an asymmetrical fashion into a closed spheroid ^6^. (B) The average number of localized point defects of each type (+1, −1/2 and +1/2) at different stages of open ring regeneration (shaded region – standard deviation). The configuration of point defects develops over time. (C) The net charge of localized point defects as a function of time in regenerating open rings. As in regeneration from excised fragments (Fig. 2D), the net charge typically reaches +2 (the topologically required value) only at ~24 hours, after the ordered regions expand and defects become localized. The defect distribution in (B,C) was determined for a population of regenerating open rings that were imaged using a spinning-disk confocal microscope from 4 directions within square FEP tubes at the specified time-points (*T=*0, 6, 24, 48, 72 hr with *N=*36, 25, 26, 27, 34 samples at each time point, respectively; see Methods). The data refers to defects that arise from the disordered regions and their evolution, and does not account for the additional defects associated with tentacle formation. (D) Images from a time-lapse spinning-disk confocal movie of a regenerating open ring. An early −1/2 defect is visible at *T*=18 hr. The −1/2 defect is retained and appears at the junction between the two body axes of the regenerated *Hydra* that has two feet and a head (*T*=50 hr).

**Figure S7.**
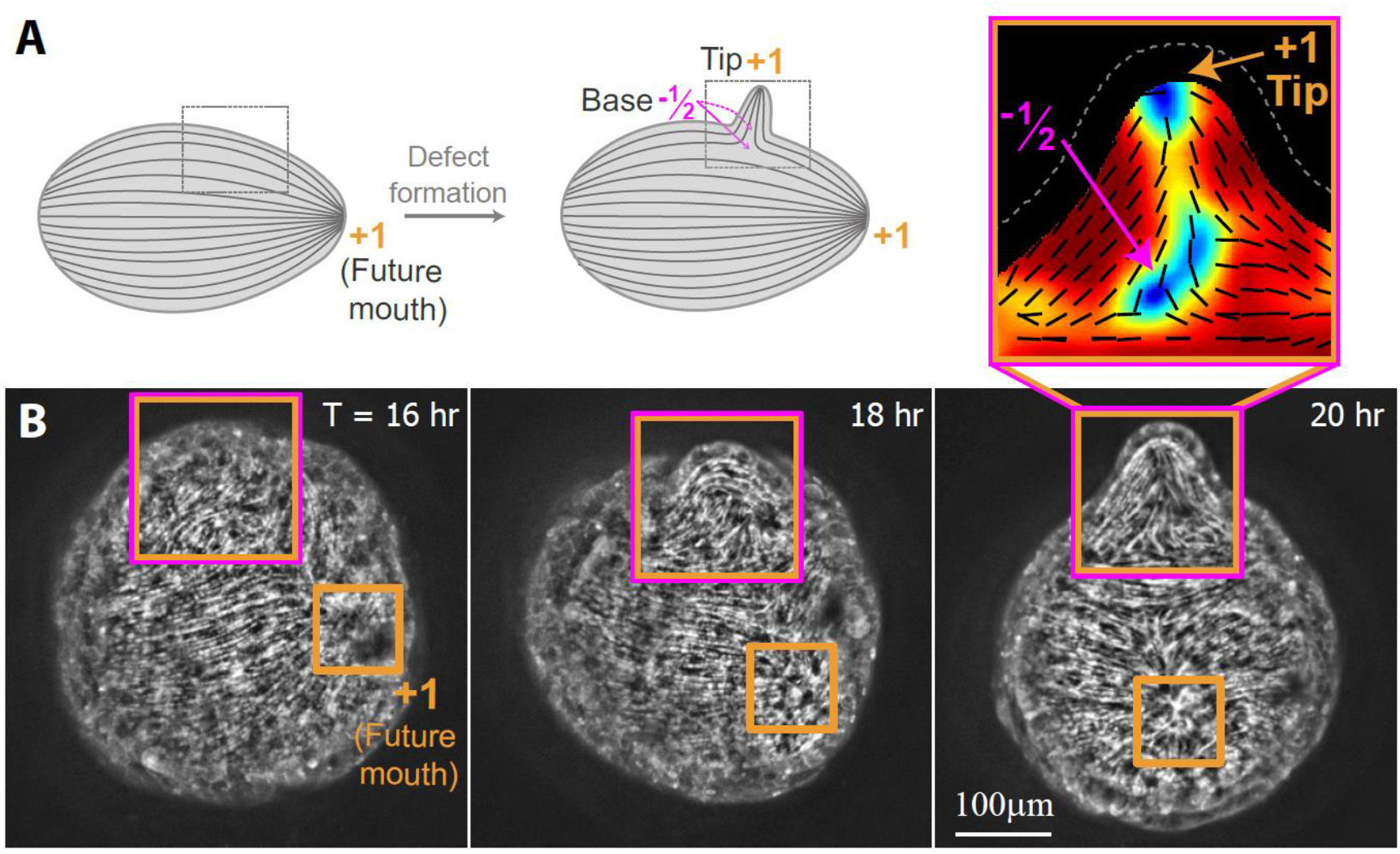
*De novo* formation of topological defects during *Hydra* regeneration. (A) Schematics of *de novo* defect formation: a +1 defect appears together with two −1/2 defects simultaneously with the formation of a local protrusion during tentacle formation. The +1 defect localizes to the region of highest positive curvature at the tip of the protrusion, whereas the two −1/2 defects localize at saddle regions of negative curvature at the base of the protrusion. The new tentacle emerges near an existing +1 defect that will become the site of the mouth in the regenerated animal. (B) Images from a time-lapse movie showing the ectodermal actin organization during the formation of a tentacle near an existing +1 defect that will become the site of the mouth of the regenerated *Hydra*. The process involves the appearance of a +1 defect at the tip of an emerging protrusion together with two −1/2 defects at opposite sides of its base, which subsequently elongates to become a tentacle in the regenerated animal. 2D projection images extracted from 3D spinning-disk confocal z-stacks of a transgenic *Hydra* expressing lifeact-GFP in the ectoderm at different time points during this process are shown. Top right: zoomed image of the protrusion showing the actin fiber orientation extracted from the image intensity gradients (black lines) and the local order parameter (color; see Methods). The +1 defect at the tip of the protrusion and the −1/2 defect on the visible side of its base are apparent.

**Figure S8.**
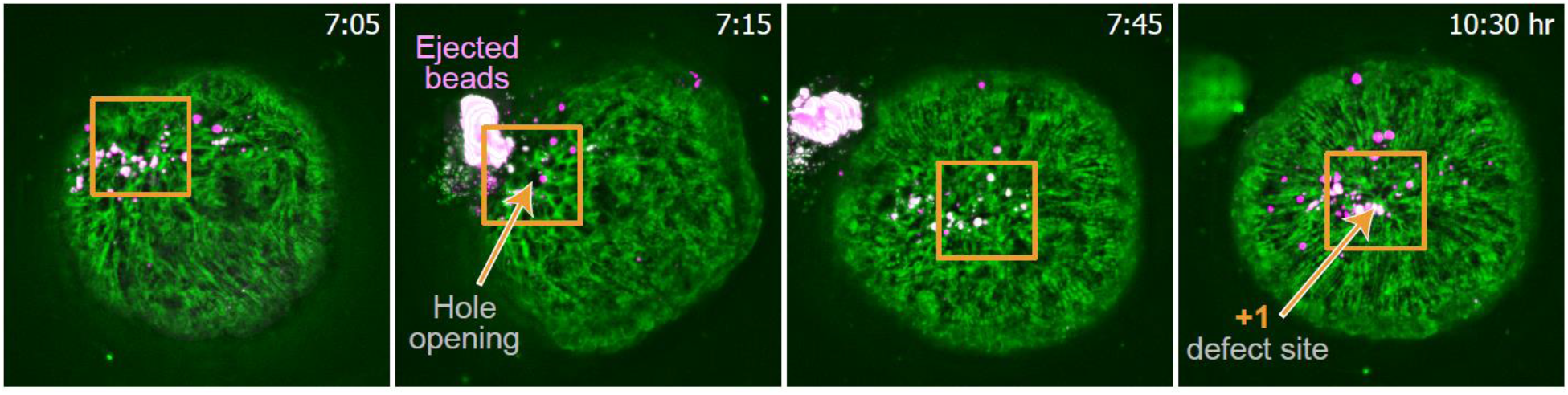
Hole formation in a disordered region prior to the appearance of a +1 defect. Images from a time-lapse movie of a regenerating fragment that was incubated with fluorescent beads (Methods). Some of the beads adhere to the tissue providing labeled landmarks on the regenerating tissue. 2D projection images extracted from 3D spinning-disk confocal z-stacks of the ectodermal lifeact-GFP (green) and the beads (magenta) are shown. Images show the sample before (at 7:05 hr:min), during (7:15), and after (7:45) the formation of a hole in a disordered region that is surrounded by regions with ordered fibers that generate a very spread out +1 defect. The beads that were trapped in the internal cavity during folding and were ejected from the hole during its rupture are visible. After several hours (10:30), following the induction of order in the region surrounding the rupture site, a localized +1 point defect develops at that site.

## Supplementary Movies

### Movie 1: Light-sheet movie of a regenerating fragment

Time-lapse light-sheet movie showing projected images of lifeact-GFP in a regenerating tissue fragment (top) and the corresponding nematic orientation field and local order parameter (bottom; Fig. 2B). Images are shown from 4 different views, each rotated by a 90-degree angle, as indicated. The movie shows the regeneration process beginning at the early tissue spheroid stage, with regions of well-aligned fibers inherited from the original *Hydra* tissue as well as disordered regions. Over the first 24-hours, the disordered regions become aligned through induction of order from neighboring ordered regions, and point defects in the nematic order develop in the previously disordered regions. The evolution of the defects can be followed to the formation of the mouth and foot defining the body axis of the regenerated animal, in addition to the later formation of tentacles. The tissue dynamics characteristic of the regeneration process, including contractions, expansions, and changes in shape are apparent. The tissue outlines are indicated with dashed lines. The images were centered to correct for movements of the whole tissue. The elapsed time from excision is displayed in hours, and the scale bar is 100 μm. The horizontal streaking of the fluorescent signal apparent in some frames is an artefact of the light-sheet illumination.

### Movie 2: Movement of a +1/2 defect relative to the underlying tissue

Time-lapse, spinning-disk confocal movie depicting the movement of a +1/2 defect relative to the underlying tissue (Fig. 3A). The +1/2 defect moves away from the labeled tissue in the direction of its rounded end. Right: images of the lifeact-GFP signal (gray scale) overlaid with the fluorescent tissue label (green). Left: corresponding nematic orientation field and local order parameter. The defect position (cyan arrow) and labeled landmark (green arrow) are indicated. The tissue outlines are indicated with dashed lines. The images were centered and rotated to correct for movements of the whole tissue. The elapsed time from excision is displayed in hours, and the scale bar is 100 μm.

### Movie 3: Annihilation of a +1/2 and a −1/2 defect

Time-lapse, spinning-disk confocal movie showing the annihilation of a +1/2 and a −1/2 defect (Fig. 3D). Near the end of the movie, the two defects (that have a net charge of 0) merge and the region becomes ordered with well-aligned parallel fibers. Right: images of the lifeact-GFP signal. Left: corresponding nematic orientation field and local order parameter. The position of the +1/2 and −1/2 defects are indicated (cyan and magenta arrows, respectively). The tissue outlines are indicated with dashed lines. The images were centered and rotated to correct for movements of the whole tissue. The elapsed time from excision is displayed in hours, and the scale bar is 100 μm.

### Movie 4: A stationary +1 defect becomes the site of mouth formation

Time-lapse, spinning-disk confocal movie showing a +1 defect that remains stationary with respect to the underlying tissue. The defect site can be followed through time and coincides with the site of head formation in the regenerated *Hydra* (Fig. 4A). The labeled cells located near the center of the +1 defect can be followed directly to the mouth in the regenerated animal. Right: images of the lifeact-GFP signal (gray scale) overlaid with the fluorescent tissue label (green). Left: corresponding nematic orientation field and local order parameter. The defect position (box) and labeled landmark (green arrow) are indicated. The tissue outlines are indicated with dashed lines. The images were centered to correct for movements of the whole tissue. The elapsed time from excision is displayed in hours, and the scale bar is 100 μm.

### Movie 5: The region between two +1/2 defects becomes the site of foot formation

Time-lapse, spinning-disk confocal movie showing two +1/2 defects with their rounded ends facing each other moving closer together. The labeled cells located between the two +1/2 defects remain centered between the defects as they move closer, and the regenerated foot forms at this site (Fig. 4B). In this case, the final merging of the two +1/2 defects into a +1 defect at the foot occurs late in the regeneration, after the establishment of the body axis. Right: images of the lifeact-GFP signal (gray scale) overlaid with the fluorescent tissue label (green). Left: corresponding nematic orientation field and local order parameter. The defect positions (cyan arrows) and labeled landmark (green arrow) are indicated when present on the visible side of the tissue. The tissue outlines are indicated with dashed lines. The images were centered to correct for movements of the whole tissue. The elapsed time from excision is displayed in hours, and the scale bar is 100 μm.

